# Deconvolving sequence features that discriminate between overlapping regulatory annotations

**DOI:** 10.1101/100511

**Authors:** Akshay Kakumanu, Silvia Velasco, Esteban Mazzoni, Shaun Mahony

## Abstract

Genomic loci with regulatory potential can be identified and annotated with various properties. For example, genomic sites may be annotated as being bound by a given transcription factor (TF) in one or more cell types. The same sites may be further labeled as being proximal or distal to known promoters. Given such a collection of labeled sites, it is natural to ask what sequence features are associated with each annotation label. However, discovering such label-specific sequence features is often confounded by overlaps between annotation labels; e.g. if regulatory sites specific to a given cell type are also more likely to be promoter-proximal, it is difficult to assess whether motifs identified in that set of sites are associated with the cell type or associated with promoters. In order to meet this challenge, we developed SeqUnwinder, a principled approach to deconvolving interpretable discriminative sequence features associated with overlapping annotation labels. We demonstrate the novel analysis abilities of SeqUnwinder using three examples. Firstly, we show SeqUnwinder’s ability to unravel sequence features associated with the dynamic binding behavior of TFs during motor neuron programming from features associated with chromatin state in the initial embryonic stem cells. Secondly, we characterize distinct sequence properties of multi-condition and cell-specific TF binding sites after controlling for uneven associations with promoter proximity. Finally, we demonstrate the scalability of SeqUnwinder to discover cell-specific sequence features from over one hundred thousand genomic loci that display DNase I hypersensitivity in one or more ENCODE cell lines.

**Availability:** https://github.com/seqcode/sequnwinder

## Introduction

Regulatory genomics analyses often focus on finding sequence features associated with genomic sites that share some property or annotation “label”. Such problems are typically phrased in terms of a classification between mutually-exclusive categories. For example, we may wish to find sequence features that discriminate between sites bound by a particular transcription factor (TF) and unbound sites, or between sites that are associated with gene activation and repression. With the increased availability of genome-wide epigenomic datasets such as those generated by the ENCODE and ROADMAP projects (ENCODE Project Consortium, 2012; Roadmap Epigenomics Consortium *et al*, 2015), it is now possible to provide a more detailed annotation of regulatory sites beyond binary labels such as “bound” and “unbound”. For example, TF binding activity can be sub-categorized according to which cell types or conditions it is bound in, or according to whether those sites display coincident ChIP-enrichment of other proteins or histone modifications. Genome segmentation methods (Ernst & Kellis, 2012; Hoffman *et al*, 2012; Zhang *et al*, 2016) provide an automated annotation of promoters, enhancers, and various other chromatin states that can also be overlaid on a TF’s binding sites. As regulatory region annotations become more complex and multi-layered, there is a growing need for computational methods that can find sequence features that discriminate between annotation labels, even if those labels overlap one another

To gain insight into the general multi-label problem that we aim to solve, consider the hypothetical scenario presented in Figure 1a, where a given TF’s binding sites have been labeled as being bound in cell types A, B, or C. The same sites are further characterized as being proximal or distal to promoters (Pr and Di, respectively), where the latter labels unevenly overlap the cell type labels. For example, let’s suppose that cell type A’s sites are more likely to be promoter proximal than sites in other cell types. If we now try to find sequence feature associated with each cell type, the uneven overlap with promoter-proximity will confound our results. It is likely that some features discovered in cell type A’s sites might actually be due to the generic properties of promoter proximal regions as opposed to being specifically associated with cell type A’s regulatory mechanisms.

**Figure 1.**
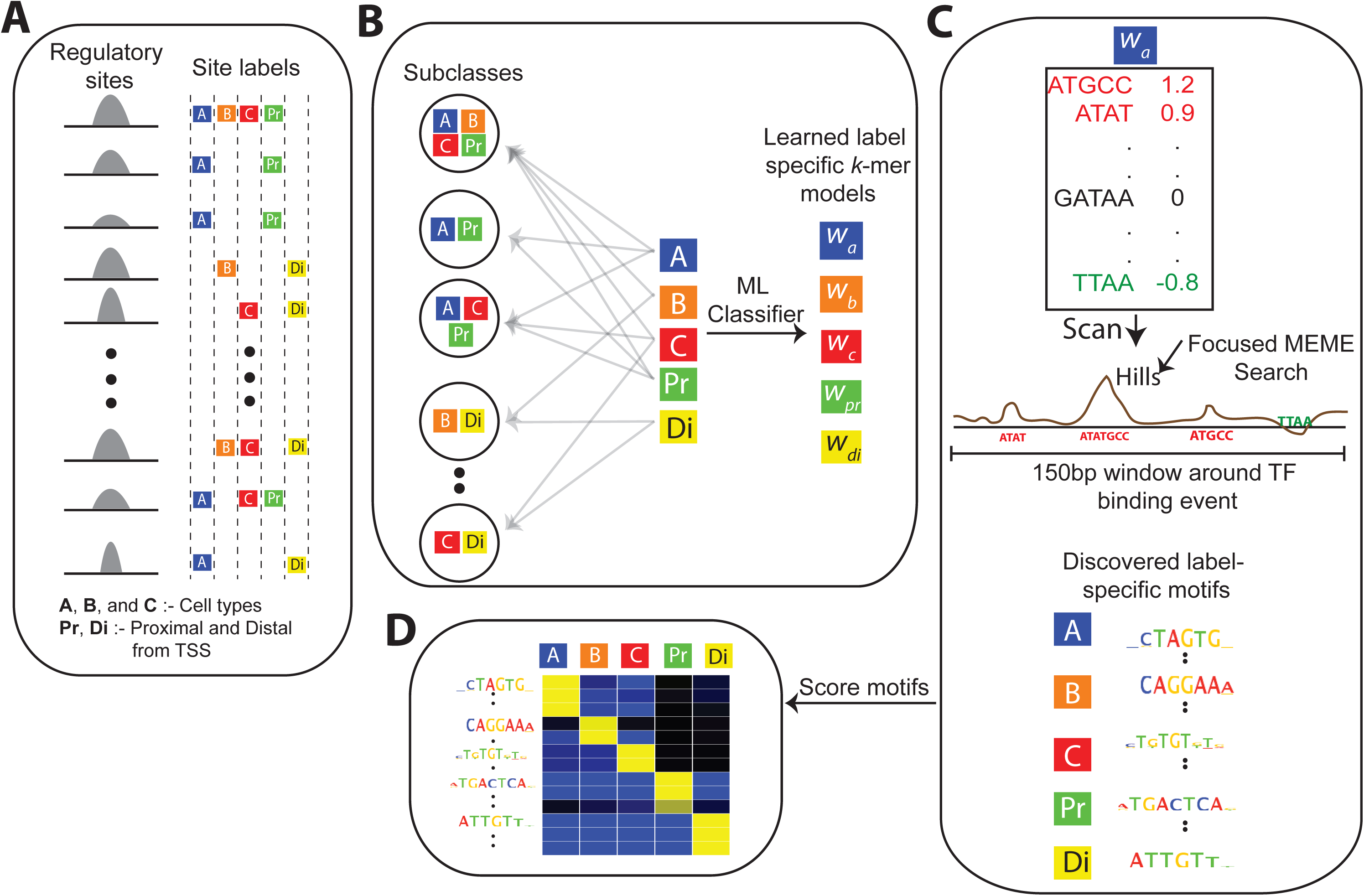
Overview of SeqUnwinder, which takes an input list of annotated genomic sites and identifies label-specific discriminative motifs. **A)** Schematic showing a typical input instance for SeqUnwinder: a list of genomic coordinates and corresponding annotation labels. **B)** The underlying classification framework implemented in SeqUnwinder. Subclasses (combination of annotation labels) are treated as different classes in a multi-class classification framework. The label-specific properties are implicitly modeled using L1-regularization. **C)** Weighted *k*-mer models are used to identify 10-15bp focus regions called hills. MEME is used to identify motifs at hills. **D)** *De novo* identified motifs in C) are scored using the weighted *k*-mer model to obtain label-specific scores.

Almost all existing classification methods for characterizing discriminative sequence features assume that the classes (i.e. annotation labels) are mutually exclusive, and therefore cannot appropriately handle scenarios such as that outlined in Figure 1a. While there have been several significant developments in characterizing discriminative sequence features using convolutional neural networks and support vector machines (SVMs) with various *k*-mer based sequence kernels (Arvey *et al*, 2012; Ghandi *et al*, 2014; Lee *et al*, 2011), these applications typically focus on discriminating between two mutually exclusive classes (Bailey, 2011; Alipanahi *et al*, 2015). Multi-class discriminative sequence feature frameworks have been proposed (Beer & Tavazoie, 2004; Elemento *et al*, 2007), but these approaches have also been limited to analysis of mutually exclusive classes.

A few existing methods do allow a limited analysis of datasets where annotation labels partially overlap, but these approaches were designed for essentially two-class classification problems where the multi-task framework enables modeling of the “common” task in addition to the two classes. For example, (Arvey *et al*, 2012) used a multi-task SVM classifier to learn cell-type specific and shared binding preferences of TF binding sites in two cell-types, and the group lasso based logistic regression classifier SeqGL (Setty & Leslie, 2015) also implements a similar multi-task framework. No existing discriminative feature discovery method is applicable to scenarios where a set of genomic sites contains multiple annotation labels with arbitrary rates of overlap between them. We therefore require a structured classification framework that can deconvolve sequence features associated with arbitrary numbers of overlapping annotation labels.

In this work, we present SeqUnwinder, a novel classification framework for characterizing interpretable sequence features associated with overlapping sets of genomic annotation labels. SeqUnwinder begins by defining genomic site subclasses based on the combinations of labels annotated at these sites (Figure 1b). The site subclasses are treated as distinct classes for a multiclass logistic regression model that uses *k*-mer frequencies as predictors. Moreover, SeqUnwinder also models each individual label’s specific features by incorporating them in an L1 regularization term (see Methods). Regularization encourages consistent features to be shared across subclasses that are spanned by a label, thus implicitly enabling label-specific features to be learned (Figure 1b). The trained classifier encapsulates weighted *k*-mer models specific to each label and each subclass (i.e. combination of labels). The label-or subclass-specific *k*-mer model is scanned across the original genomic sites to identify focused regions (which we term “hills”) that contain discriminative sequence signals (Figure 1c). Finally, to aid interpretability, SeqUnwinder identifies over-represented motifs in the hills and scores them using label-and subclass-specific *k*-mer models (Figure 1d).

We demonstrate the unique analysis abilities of SeqUnwinder using both synthetic sequence datasets and collections of real TF ChIP-seq and DNase-seq experiments. In the real datasets, we begin with a motivating example that analyzes TF binding during the programming of embryonic stem (ES) cells into induced motor neurons (iMNs) (Mazzoni *et al*, 2013; Velasco *et al*, 2017). In this example, we categorize the TF binding sites according to dynamic binding behaviors observed during the programming process. These binding site categories are further (unevenly) split into subclasses according to whether they are in an accessible/active chromatin state in the initial ES cells. We demonstrate that SeqUnwinder can deconvolve sequence features associated with binding dynamics from those associated with initial chromatin state, thereby providing testable hypotheses about the binding mechanisms associated with each annotation label.

In two further examples using real epigenomic datasets, we characterize sequence features associated with genomic locations that display regulatory properties across multiple cell types. Sites that are bound by a particular TF in multiple cell types (i.e. “shared” or multi-condition sites) are often strongly biased towards being located in gene promoter regions, in contrast to cell-specific binding sites, which are typically distally located (Yip *et al*, 2012; Wang *et al*, 2012; Kheradpour & Kellis, 2014). After controlling for such biases by incorporating labels that annotate proximal and distal sites, SeqUnwinder discovers that shared TF binding sites are characterized by stronger instances of the cognate binding motif than are present at cell-specific sites. In our final example, we demonstrate that SeqUnwinder scales naturally to analyses of over one hundred thousand sites annotated with dozens of label combinations. To show this, we characterize the sequence features at shared and cell-type specific DNase I hypersensitive sites in six different ENCODE cell lines. Interestingly, we find that motifs enriched in cell-type specific DNase I hypersensitive sites are also highly enriched at cell-type specific TF binding sites for a majority of the examined TFs.

In summary, SeqUnwinder can analyze large collections of genomic loci that have been annotated with several overlapping labels to extract interpretable sequence features associated with each label.

## Results

### SeqUnwinder deconvolves sequence features associated with overlapping labels

To demonstrate the properties of SeqUnwinder, we simulated 9,000 regulatory regions and annotated each of them with labels from two overlapping sets: A, B, C and X, Y (Figure 2a). We assigned a different motif to each label. At 70% of the sites associated with each label, we inserted appropriate motif instances by sampling from the distributions defined by the position-specific scoring matrices of label assigned motifs (Figure 2a). We used this collection of sequences and label assignments to compare SeqUnwinder with a simple multi-class classification approach (MCC). In MCC training, each label was treated as a distinct class and therefore each regulatory site is included multiple times in accordance with its annotated labels.

**Figure 2.**
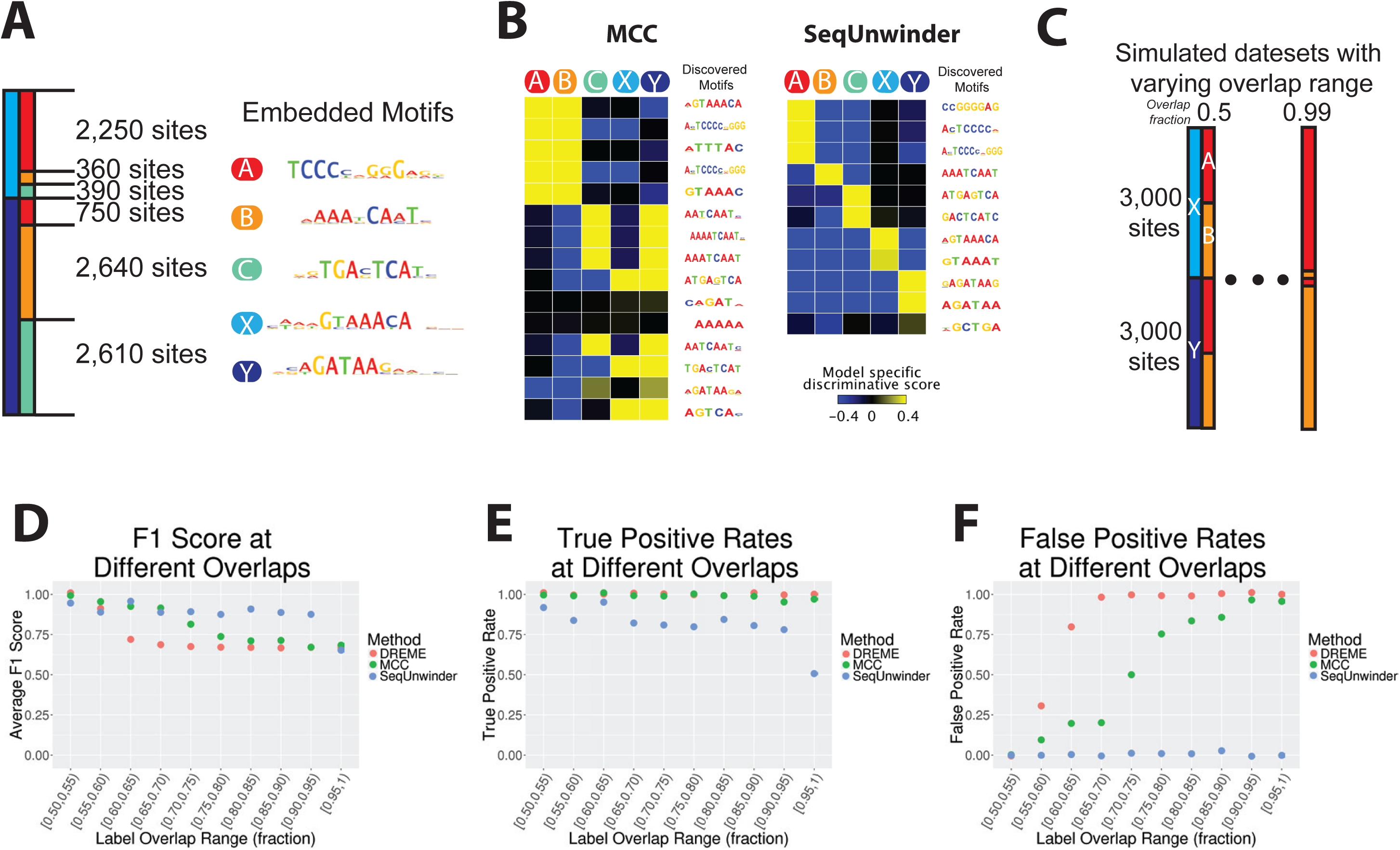
Performance of SeqUnwinder on simulated datasets. **A)** 9000 simulated genomic sites with corresponding motif associations. **B)** Label-specific scores for all *de novo* motifs identified using MCC (left) and SeqUnwinder (right) models on simulated genomic sites in “A”. **C)** Schematic showing 100 genomic datasets with 6000 genomic sites and varying degrees of label overlap ranging from 0.5 to 0.99. **D)** Performance of MCC (multi-class logistic classifier), DREME, and SeqUnwinder on simulated datasets in “C”, measured using the F1-score, **E)** true positive rates, and **F)** false positive rates.

SeqUnwinder and MCC models correctly identify motifs similar to all inserted motifs (Figure 2b). However, the MCC approach makes several incorrect motif-label associations potentially due to high overlap between labels. On the other hand, the label specific scores of the identified motifs in the SeqUnwinder model are not confounded by overlap between annotation labels. For example, even though labels X and A highly overlap, SeqUnwinder correctly assigns each motif to its respective label.

Next, we assessed the performance of SeqUnwinder at different levels of label overlaps. We simulated 100 datasets with 6000 simulated sequences, varying the degree of overlap between two sets of labels ({A, B} and {X, Y}) from 50% to 99% (Figure 2c). We then compared SeqUnwinder with MCC and DREME (Bailey, 2011), a popular discriminative motif discovery tool. Since DREME takes only two classes as input: a foreground set and a background set, we ran four different DREME runs for each of the four labels. We calculated the true positive (discovered motif correctly assigned to a label) and false positive (discovered motif incorrectly assigned to a label) rates based on the true (simulated) label assignments. We used these measures to calculate the F1 score (harmonic mean of precision and recall) at different overlapping levels (Figure 2d).

Figure 2d demonstrates the range of label overlap rates in which SeqUnwinder outperforms the alternative approaches. When the labels are uncorrelated (i.e. low or random overlap), the sequence features associated with each label do not confound one another and thus all methods perform similarly well in characterizing label-specific motifs. On the other hand, when the labels are highly correlated (i.e. high overlap), it becomes impossible for any method to correctly assign sequence features to the correct labels. SeqUnwinder performs better than the other approaches in the intermediate range of label overlaps, and accurately characterizes label-specific sequence features even when the simulated labels overlap at 90% of sites. More specifically, SeqUnwinder consistently has a false positive rate (incorrectly assigning motifs to labels) of zero at the cost of a modest decrease in true positive rates (recovering all motifs assigned to a label) (Figure 2e and 2f)

Taken together, the synthetic data experiments demonstrate that SeqUnwinder provides the ability to discover sequence features associated with highly overlapping sets of genomic site labels, outperforming other available methods.

### SeqUnwinder uncovers co-factor driven TF binding dynamics during iMN programming

To demonstrate its unique abilities in a real analysis problem, we use SeqUnwinder to study TF binding during induced motor neuron (iMN) programming. Ectopic expression of Ngn2, Isl1, and Lhx3 in mouse ES cells efficiently converts the resident ES cells into functional spinal motor neurons (Mazzoni *et al*, 2013; Velasco *et al*, 2017). We recently characterized the dynamics of motor neuron programming by studying TF binding, chromatin dynamics, and gene expression over the course of the 48hr programming process (Velasco *et al*, 2017). We found that two of the ectopically expressed TFs, Isl1 & Lhx3, bind together at the vast majority of their targets during the programming process. We also found that this cooperative pair of TFs shifted their binding targets during programming, and we used three mutually exclusive labels – early, shared, and late – to annotate Isl1/Lhx3 binding sites according to their observed dynamic occupancy patterns. Early sites were bound by Isl1/Lhx3 only during earlier stages of programming, shared sites were constantly bound over the entire programming process, and late sites were only bound during the final stage of programming.

In our previous work, we demonstrated that the early Isl1/Lhx3 sites were more accessible in the initial pluripotent cells, and we suggested that some early sites are the result of opportunistic Isl1/Lhx3 binding to ES enhancer regions (Velasco *et al*, 2017). However, this raises a question: if we discover sequence features at early sites, how can we tell if those features are specifically associated with Isl1/Lhx3 as opposed to reflecting on coincident properties of ES enhancers? In order to assess the potential confounding effects of ES regulatory sites, we trained a random forest classifier to further categorize all Isl1/Lhx3 bound sites using two additional labels: “ES-active and “ES-inactive” (see Methods). Annotating Isl1/Lhx3 sites using both sets of labels (Isl1/Lhx3 binding dynamics and ES activity) results in six different subclasses. As can be seen from Figure 3a, early sites have a higher propensity to also be active prior to ectopic TF expression in the starting ES cells. Conversely, the late sites are more likely to be inactive in ES cells.

**Figure 3.**
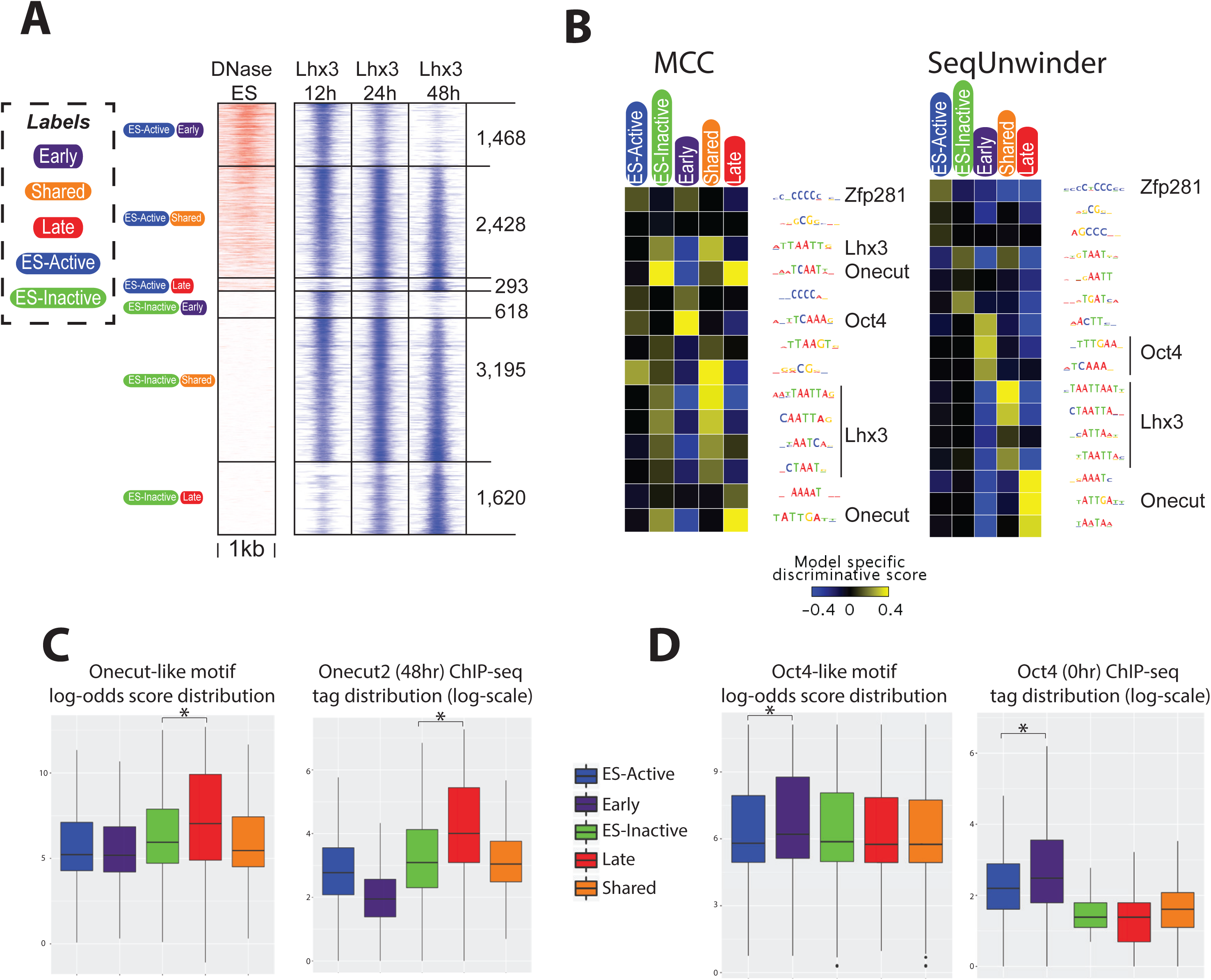
SeqUnwinder analysis of Lhx3 binding classes during iMN programming. **A)** Lhx3 binding sites labeled using their dynamic binding behavior and ES chromatin activity statuses. **B)** Label-specific scores of *de novo* motifs identified at Lhx3 binding sites defined in “A” using MCC (left) and SeqUnwinder (right) models. **C)** Distribution of Onecut2 (48hr) ChIP-seq tag counts in log-scale at “ES-active”, “ES-inactive“, “Early”, “Shared”, and “Late” sites (left panel). Log-odds score distribution of *de novo* discovered Onecut-like motif at “ES-active”, “ES-inactive“, “Early”, “Shared”, and “Late” sites (right panel). **D)** Distribution of Oct4 (0hr) ChIP-seq tag counts in log-scale at “ES-active”, “ES-inactive“, “Early”, “Shared”, and “Late” sites (left panel). Log-odds score distribution of *de novo* discovered Oct4-like motif at “ES-active”, “ES-inactive“, “Early”, “Shared”, and “Late” sites (right panel). Statistical significance calculated using Mann-Whitney-Wilcoxon test (*: P-value < 0.001)

We next trained SeqUnwinder on the multi-label Isl1/Lhx3 dataset, and compared the results with those of the simple MCC approach described in the previous section (Figure 3b). Both the MCC approach and SeqUnwinder discover similar sets of motifs. For example, both approaches find motifs corresponding to the binding preferences of Oct4, Zfp281, Onecut-family TFs, and homeobox motifs corresponding to the cognate Isl1/Lhx3 binding preference (Figure 3b). However, the two approaches produce different associations between motifs and labels. For example, the MCC approach associates the Oct4 motif with both the “early” and “ES-active” labels, and it associates the Onecut motif with both “late” and “ES-inactive” labels (Figure S1). SeqUnwinder, on the other hand, makes much cleaner associations; the Oct4 motif is only associated with the “early” label, and the Onecut motif is only associated with the “late” label. These preferential SeqUnwinder associations suggest that Oct4 and Onecut motifs are not merely coincidental features due to the ES activity status of the binding sites.

The SeqUnwinder motif-label associations suggest that Isl1/Lhx3 bind cooperatively with Oct4 and Onecut TFs at subsets of early and late binding sites, respectively. As described in our earlier work, we characterized Onecut2 binding to be highly enriched at late Isl1/Lhx3 sites during iMN programming (Velasco *et al*, 2017). We also found that late sites are not bound by Isl1/Lhx3 (and iMN programming does not proceed) in cellular conditions under which Onecut TFs are not expressed (Velasco *et al*, 2017), supporting a model in which late Isl1/Lhx3 binding is dependent on Onecut TFs. Analysis of motif log-odds scores and Onecut2 ChIP enrichment further support SeqUnwinder’s prediction that the Onecut motif is not merely a general feature of ES-inactive sites (Figure 3c).

Conversely, Oct4 is predicted by SeqUnwinder to be a specific feature of “early” binding sites, and not merely an artifact associated with “ES-active” sites. Using ChIP-seq, we profiled the binding of Oct4 immediately before NIL induction. As shown in Figure 3d, Oct4 motif log-odds scores and ChIP-seq tags show a preferential enrichment at early Isl1/Lhx3 sites, in line with SeqUnwinder’s prediction.

Interestingly, the motif features that are most highly associated with shared binding sites all correspond to homeobox motifs of the type bound by Isl1/Lhx3 (Figure 3b and Figure S1). One possible explanation is that there are stronger or more frequent cognate motif instances at sites bound by a given TF across multiple timepoints, or indeed across multiple unrelated cell types. We further assess this hypothesis in the following section.

Our analysis of Isl1/Lhx3 binding during iMN programming serves as an example analysis scenario in which SeqUnwinder identifies motif features associated with multiple overlapping labels, leading to testable hypotheses about co-factors that serve mechanistic roles at subsets of binding sites. Specifically, SeqUnwinder’s analysis of the iMN programming system suggests that Oct4 and Onecut motifs are not merely generic features of the starting and finishing cell types, and instead these TFs may specifically collaborate with Isl1/Lhx3 at different stages in the programming process.

### Multi-condition TF binding sites are characterized by stronger cognate motif instances

The sequence properties of tissue-specific TF binding sites have been extensively studied (Heinz *et al*, 2010; Arvey *et al*, 2012; Setty & Leslie, 2015). As might be expected, sites that are bound by a given TF in only one cell type are often enriched for motifs of other TFs expressed in that cell type. Therefore, a given TF’s cell-specific binding activity is likely determined by context-specific interactions with other expressed regulators.

Most TFs also display cell-invariant binding activities. In other words, each TF typically has a cohort of sites that appear bound in all or most cellular conditions in which that TF is active. Despite the potential regulatory significance of such multi-condition binding sites, little is known about the sequence properties that enable a TF to bind them regardless of cellular conditions. Studies of individual TFs suggest that binding affinity to cognate motif instances may play a role in distinguishing multi-condition binding sites from tissue-specific sites (Gertz *et al*, 2013; Mahony *et al*, 2014).

In order to characterize sequence discriminants of multi-condition TF binding sites across a wider range of TFs, we curated multi-condition ChIP-seq experiments from the ENCODE project. We restricted our analysis to the 17 sequence-specific TFs profiled in all 3 primary ENCODE cell-lines (K562, GM12878, and H1-hESC; see Methods) (ENCODE Project Consortium, 2012). For each TF, we curated sets of tissue-specific sites in each cell type, and a further set of sites that are “shared” across all three cell types (see Methods). For most examined TFs, the majority of shared binding sites were located in promoter proximal regions (Figure S2). Promoter proximal sites are known to have distinct sequence biases, which could confound the discovery of sequence features associated with shared sites. We therefore further labeled each TF’s binding sites as being located proximal or distal to annotated TSSs. Introducing the proximal and distal labels should marginalize the proximal bias at shared sites, as the sequence features learned by SeqUnwinder at shared sites must be consistently enriched at both proximal and distal sites. In summary, each examined TF’s binding sites is categorized into eight subclasses, each of which is composed of combinations of six distinct labels.

We applied SeqUnwinder to each labeled sequence collection in order to characterize label-specific sequence features. We illustrate the process with SeqUnwinder’s results for YY1. We started with a total of ∼35,000 YY1 binding events called by MultiGPS across the three cell types, categorized into the eight aforementioned subclasses (Figure 4a). SeqUnwinder identifies several *de novo* motifs in this collection (Figure 4b). Interestingly, SeqUnwinder predicts that a motif matching the cognate YY1 motif is strongly associated with the “shared” label. The cell-type specific, proximal and distal labels had low or negative scores for this cognate motif. Note here that a non-positive label-specific score for a motif does not necessarily imply complete absence of instances of that motif. A significant depletion of motif instances at sites annotated by a label compared to other labels can very likely result in non-positive scores. Cell-type specific sites had higher scores for co-factor motifs. For example, H1-hESC specific sites were enriched in instances of a TEAD-like motif, while K562-specific sites and GM12878-specific sites were enriched for a GATA-like motif and an IRF-like motif, respectively. In fact, GATA2 ChIP-seq reads in K562, IRF4 ChIP-seq reads in GM12878, and TEAD4 ChIP-seq reads in H1hESC showed striking enrichment at corresponding cell-specific YY1 binding sites (Figure 4a).

**Figure 4.**
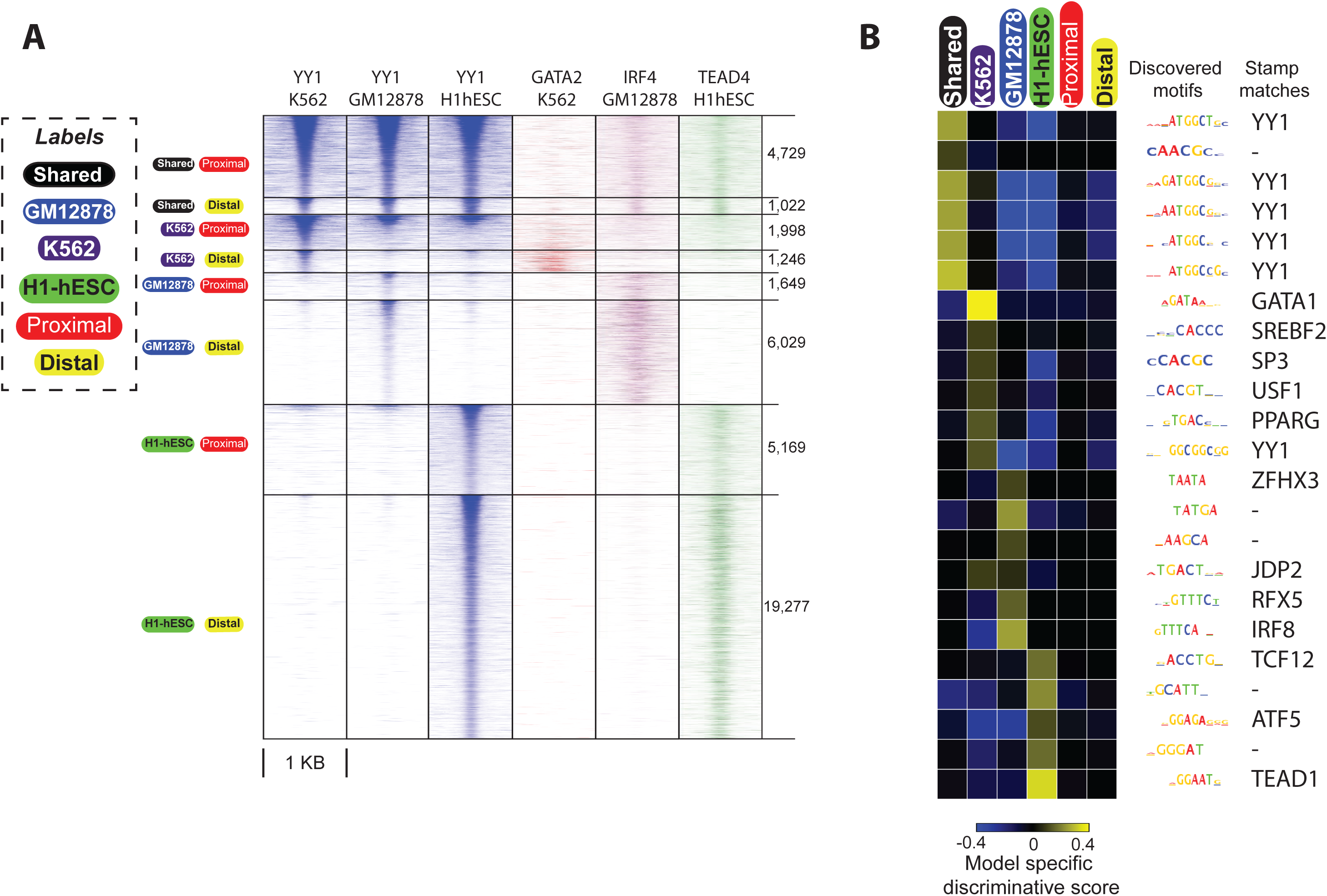
SeqUnwinder analysis of sequence features at multi-condition TF binding sites for ENCODE YY1 datasets. **A)** Heatmaps showing the YY1 ChIP-seq reads at curated YY1 binding sites, stratified based on binding across cell-lines and distance from annotated mRNA TSS. The order of subclasses is: Shared and Proximal, Shared and Distal, K562 and Proximal, K562 and Distal, GM12878 and Proximal, GM12878 and Distal, H1-hESC and Proximal, and H1-hESC and Distal. **B)** *De novo* motifs and corresponding label specific scores identified using SeqUnwinder at events defined in A).

Analogous results were observed for many of the examined factors. For 13 out of the 17 examined factors, SeqUnwinder predicts that the cognate motif is highly associated with the “shared” label (Figure 5a). Despite significant overlaps between shared sites and promoter proximal sites (Figure S2), the cognate motifs were not found to be predictive of any TF’s “proximal” label (Figure 5a). Further, the cognate motif was not specifically predictive of cell-type-specific labels for any of the examined TFs, with the exception of H1-hESC-specific sites for CEBPB, NRSF and SRF. An orthogonal analysis of log-odds motif scoring distributions across each TF’s labels is consistent with the SeqUnwinder results (Figure S3). Thus, after implicitly controlling for uneven overlaps with promoter regions, SeqUnwinder predicts that higher affinity or more frequent cognate motif instances are a common feature of multi-condition TF binding sites.

**Figure 5.**
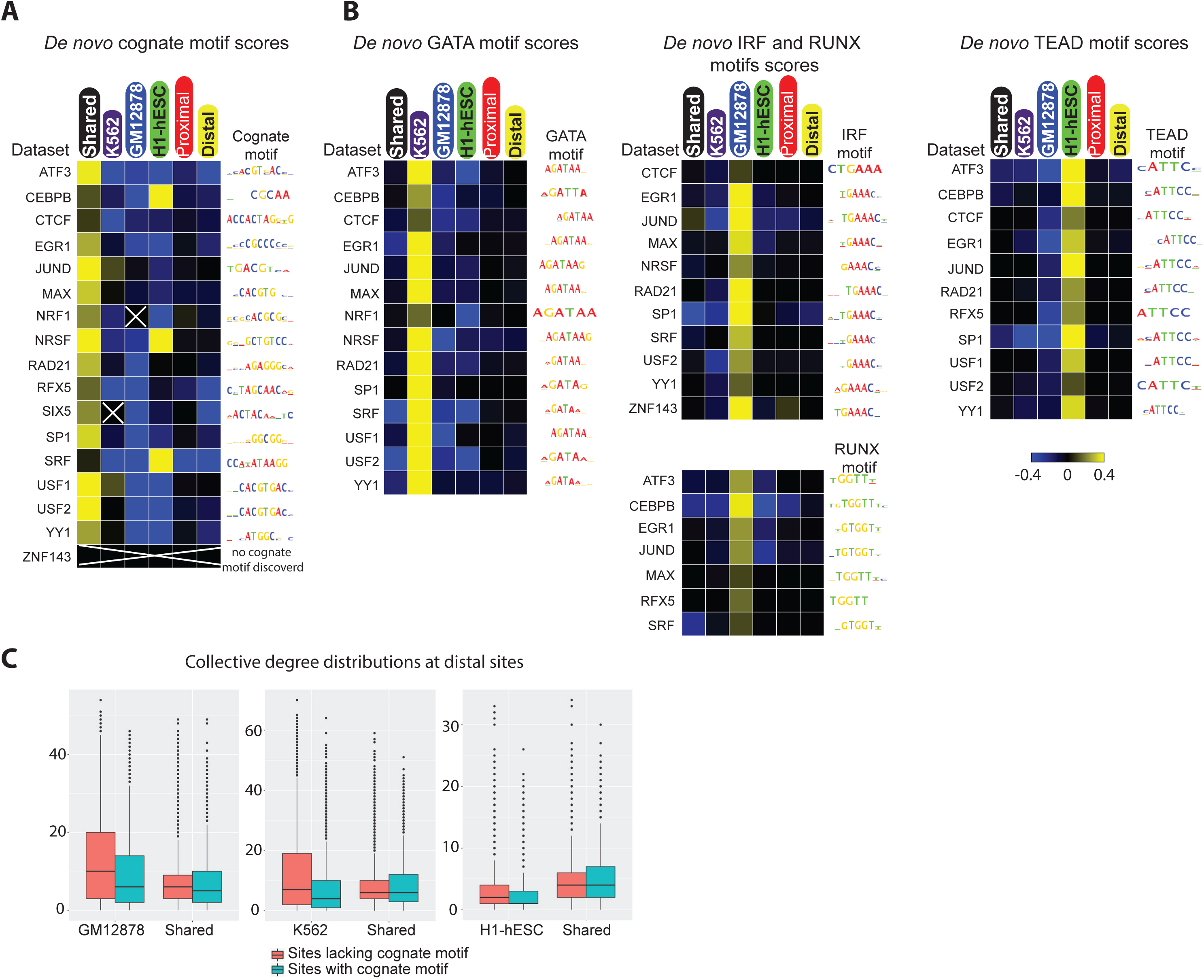
SeqUnwinder analysis of sequence features at multi-condition TF binding sites for 17 ENCODE TFs. **A)** Label specific scores of *de novo* discovered cognate motifs across all 17 ENCODE TF datasets. SeqUnwinder did not discover a cognate motif for ZNF143. GM12878-enriched sites for NRF1 and H1-hESC-specific sites for SRF were excluded because of low number of binding sites. **B)** Label specific scores of *de novo* discovered GATA-like, IRF-like, RUNX-like and TEAD-like motifs. **C)** Collective degree distributions at distal shared and cell-specific sites further stratified based on presence of cognate motif.

We also examined the motifs that SeqUnwinder predicts to be associated with cell-type-specific binding labels. Interestingly, we found IRF and RUNX motifs enriched at GM12878-specific binding sites for 11 and 7 of the 17 examined TFs, respectively. Similarly, the GATA motif was predictive of K562-specific binding for 14 out of the 17 examined TFs. A TEAD-like motif was predictive of H1-hESC specific sites for 11 of the 17 TFs (Figure 5b).

As demonstrated, SeqUnwinder predicts that cell-type-specific sites are depleted for cognate motif instances but are enriched for motif instances of other lineage-specific regulators. These results are consistent with the “TF collective” model proposed by Junion and colleagues (Junion *et al*, 2012). Under this model, the cooperative binding of large numbers of TFs is driven by the presence of motifs for a subset of lineage-specific factors that drive recruitment of the rest (i.e. the motifs for some TFs need not always be present).

To further support the “TF collective” interpretation of SeqUnwinder’s results, we tested the degree to which TSS-distal cell-type-specific sites are bound by numerous other TFs. We first defined a binding site’s “collective degree” as the number of distinct TFs with nearby ChIP-seq peaks. To calculate collective degree, we used a total of 158, 102, and 202 ChIP-seq datasets in GM12878, H1-hESC, and K562 cell-types, respectively. From Figure 5c, it is clear that distal K562-and GM12878-specific sites lacking a cognate motif instance have higher collective degrees. Similar findings were previously identified at the “high occupancy of transcription-related factors (HOT)” regions (Yip *et al*, 2012).

In summary, our analyses demonstrate the applicability of SeqUnwinder to the increasingly common problem of characterizing sequence features associated with cell-specific and cell-invariant TF binding. Uniquely, SeqUnwinder can implicitly account for location-dependent sequence composition biases by incorporating knowledge of extra layers of annotations. Our results support the model that high affinity cognate motif instances are a striking feature of multi-conditionally bound sites across a broad range of TFs (Gertz *et al*, 2013; Mahony *et al*, 2014). Furthermore, our results support a model in which cell-type-specific sites lacking cognate motif instances are bound in a “TF collective” fashion.

### SeqUnwinder identifies sequence features at shared and cell-specific DHS in six different ENCODE cell-lines

Finally, we aim to demonstrate the utility of SeqUnwinder in identifying sequence features at large numbers of genomic loci annotated with several labels. We first annotated a large collection of DNase I hypersensitive (DHS) sites with six cell-line labels depending on the enrichment of DNase-seq reads (Figure 6a). If we had used analysis methods that rely on mutually exclusive categories, we would need to restrict analysis to ∼97,000 sites labeled as either shared or exclusive to one of the six cell types (Shen *et al*, 2012). Indeed, these strict category definitions may introduce sequence composition biases into each category. However, by taking advantage of SeqUnwinder’s unique framework to pool information from all subclasses, we can analyze ∼140,000 DHS sites that we annotate into 22 subclasses as shared (i.e. enriched in 5 or more cell types) or specific to one or two cell types (Figure 6a).

**Figure 6.**
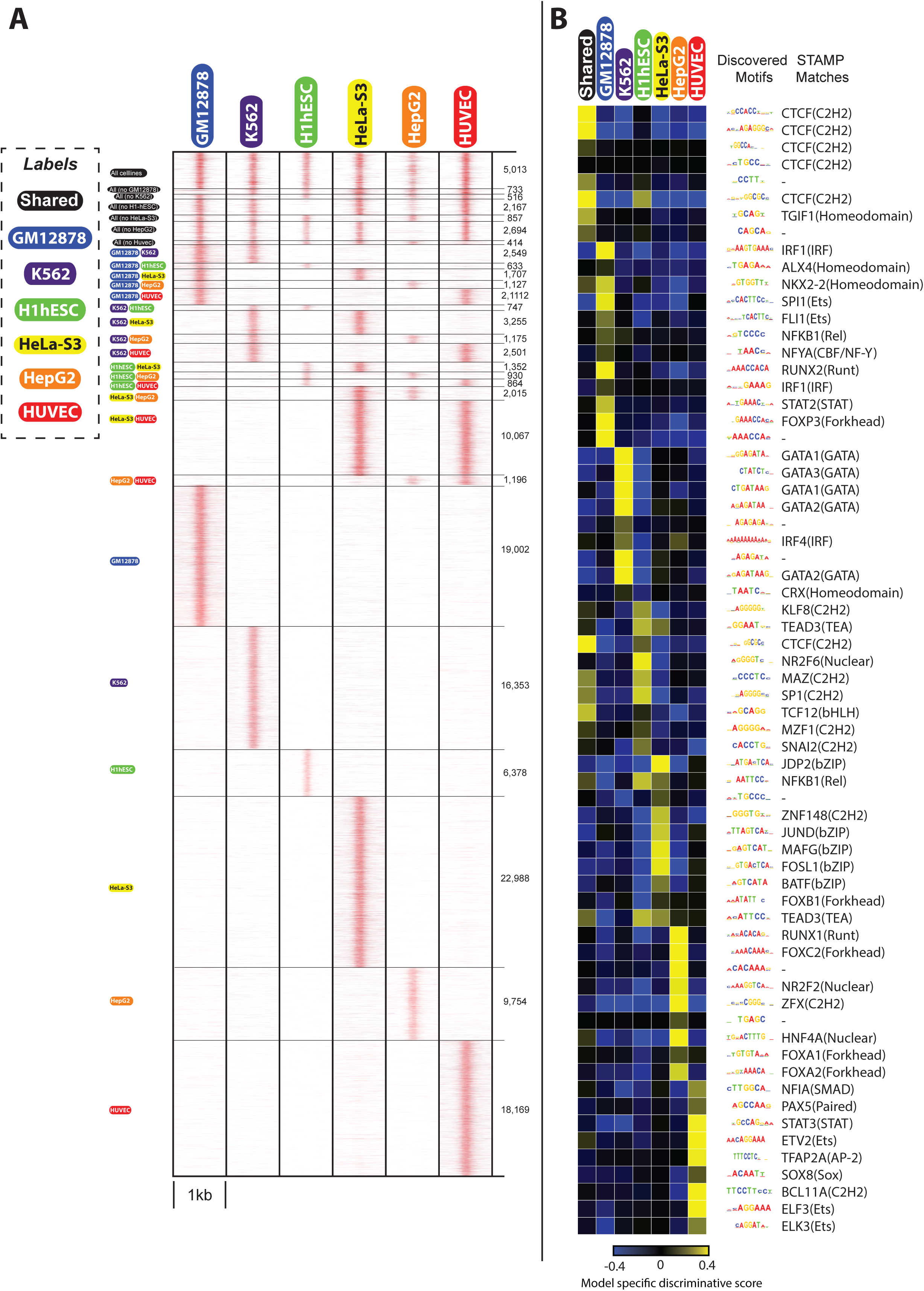
Discriminative sequence feature analysis at DHS sites in 6 different ENCODE cell-lines using SeqUnwinder. **A)** ∼140K DHSs sites annotated with 6 different cell-line labels used to identify cell-line specific and shared sequence features. **B)** Label specific scores of all the *de novo* motifs identified at DHSs sites in “A”.

SeqUnwinder identifies several motifs in this large collection of DHS sites, including those previously associated with specific cell-types (Wang *et al*, 2012; Kheradpour & Kellis, 2014; Lu *et al*, 2017) (Figure 6b). For example, components of the CTCF motif were highly predictive of shared DHS sites. This result is consistent with previous findings suggesting relatively invariant CTCF binding across cellular contexts (Cuddapah *et al*, 2009; Kim *et al*, 2007). RUNX, IRF and NF-κB motifs were enriched at GM12878-specific DHS sites. These motifs were also discovered by others at GM12878-specific DHS sites (Kheradpour & Kellis, 2014; Setty & Leslie, 2015). Motifs corresponding to GATA TFs, key regulators of erythroid development (Pevny *et al*, 1995; Welch *et al*, 2004; Han *et al*, 2016), were enriched at K562-specific DHS sites. SNAI and TEAD motifs were enriched at H1-hESC sites, consistent with previous observations (Setty & Leslie, 2015). JUND and FOS motifs were enriched at HeLa-S3-specific DHS sites. Motifs for HNF4A and FOX TFs, known master regulator of hepatocytes (Duncan *et al*, 1998; Friedman & Kaestner, 2006; DeLaForest *et al*, 2011; Alder *et al*, 2014), were enriched at HepG2-specific DHS sites. Finally, motifs belonging to the ETS class of TFs were enriched at HUVEC-specific DHS sites (Figure 6b). ETS factors have been shown to directly convert human fibroblasts to endothelial cells (Morita *et al*, 2015). Interestingly, some of the motifs associated with cell-type specific DHS sites were also found in our analyses of cell-type specific TF binding sites above (Figure 5b). For example, IRF, GATA, and TEAD motifs associated with GM12878, K562, and H1-hESC specific DHSs were also predictive of corresponding cell-type specific binding for many of the analyzed TFs.

These results demonstrate that SeqUnwinder scales effectively in characterizing sequence features at thousands of regulatory regions annotated by several different overlapping labels.

## Discussion

Classification models have shown great potential in identifying sequence features at defined genomic sites. For example (Lee *et al*, 2011), trained an SVM classifier to discriminate putative enhancers from random sequences using an unbiased set of *k*-mers as predictors. The choice of kernel function is key to the performance of an SVM classifier. Several variants of the basic string kernel (e.g. mismatch kernel (Leslie & Kuang, 2004), di-mismatch kernel (Arvey *et al*, 2012), wild-card kernel (Leslie & Kuang, 2004; Setty & Leslie, 2015), and gkm-kernel (Ghandi *et al*, 2014)) have been proposed and have been shown to substantially improve the classifier performance. Several complementary methods using DNA shape features in a classification framework have also provided insight on the role of subtle shape features that distinguish bound from unbound sites (Zhou *et al*, 2015; Chiu *et al*, 2016; Mathelier *et al*, 2016). More recently, deep learning models have also been harnessed to predict TF binding sites from unbound sites (Alipanahi *et al*, 2015).

In this manuscript, we focus not on the form of the training features, but rather on the tangential problem of identifying sequence features that discriminate several annotations applied to a set of genomic locations. Most existing methods have been developed and optimized to identify sequence features that discriminate between mutually exclusive classes (e.g. bound and unbound sites). However, when considering different sets of genomic annotation labels, overlaps between them are very likely and can confound results. To systematically address this, we developed SeqUnwinder.

We have shown that SeqUnwinder provides a unique ability to deconvolve discriminative sequence features at overlapping sets of labeled sites. SeqUnwinder leverages overlaps between labels and identifies features that are consistently shared across subclasses spanned by a label. SeqUnwinder is easy to use and takes as input a list of genomic coordinates with corresponding annotations and identifies interpretable sequence features that are enriched at a given label or at combinations of labels (subclasses). SeqUnwinder implements a multi-threaded version of the ADMM (Boyd *et al*, 2011) framework to train the model and typically runs in less than a few hours for most datasets. SeqUnwinder can also be easily extended to incorporate different kinds of kernels and shape features.

We demonstrated the unique analysis abilities of SeqUnwinder using three analysis scenarios based on real ChIP-seq and DNase-seq datasets. Our applications are chosen to demonstrate that SeqUnwinder has the ability to predict the identities of TFs responsible for particular regulatory site properties, while accounting for potential sources of bias.

For example, in our previous characterization of Isl1/Lhx3 binding dynamics during motor neuron programming, we discovered motifs that were enriched at early and late binding site subsets (Velasco *et al*, 2017). However, our analyses were potentially confounded by a correlation between TF binding dynamics and the chromatin properties of the sites in the pre-existing ES cells. Therefore, the motifs that we previously assigned to early or late TF binding behaviors could have been merely associated with ES-active and ES-inactive sites, respectively. By implicitly accounting for the effects of overlapping annotation labels, SeqUnwinder can deconvolve sequence features associated with motor neuron programming dynamics and ES chromatin status. Our analyses support an association between Oct4 binding and early Isl1/Lhx3 binding sites, along with our previously confirmed association between Onecut TFs and late Isl1/Lhx3 binding sites (Velasco *et al*, 2017).

Our analyses of ENCODE ChIP-seq and DNase-seq datasets demonstrate the flexibility and scalability of SeqUnwinder. In analyzing TF binding across multiple cell types, we used SeqUnwinder to account for promoter proximity as a potential confounding feature. Our results add to the growing evidence that multi-condition TF binding sites tend to be distinguished by better quality instances of the primary cognate motif. For example, Gertz *et al.*, showed that ER (estrogen receptor) binding sites bound in both ECC1 and T4D7, two human cancer cell lines, had high affinity instances of EREs (estrogen response elements) compared to cell-specific binding sites. Indeed, even the “shared” binding sites for Isl1/Lhx3 in our first demonstration are characterized by stronger instances of the Isl1/Lhx3 cognate binding motifs (Figure 3b). These results suggest that many TFs have a set of binding sites that are bound across a broad range of cellular contexts, and which are characterized by better quality cognate motif instances.

Interestingly, SeqUnwinder discovers consistent motif features to be predictive of cell-specific binding sites across several examined TF ChIP-seq collections. For example, SeqUnwinder discovers GATA, IRF and TEAD motifs at K562-, GM12878-and H1hESC-specific TF binding sites, respectively. These same motifs are also discovered by SeqUnwinder to be predictive of appropriate cell-specific DNase I hypersensitivity in a large collection of DHS sites across 6 different cell types. SeqUnwinder’s characterization of cell-specific motif features in collections of DNase-seq datasets may therefore serve as a source of predictive features for efforts that aim to predict cell-specific TF binding from accessibility experimental data alone (Pique-Regi *et al*, 2011; Kähärä & Lähdesmäki, 2015; Mathelier *et al*, 2016).

In summary, SeqUnwinder provides a flexible framework for analyzing sequence features in collections of related regulatory genomic experiments, and uniquely enables the principled discovery of sequence motifs associated with multiple overlapping annotation labels.

## Methods

### SeqUnwinder model

The core of SeqUnwinder is a multiclass logistic regression classifier trained on subclasses of genomic sites. The predictive features for the classifier are *k*-mer frequencies in a fixed window around input loci, with *k* usually ranging from 4 to 6. The parameters of SeqUnwinder are *k*-mer weights for each subclass (combination of annotation labels). In addition, SeqUnwinder also models the label-specific *k*-mer weights by incorporating them in the L1 regularization term. Briefly, label-specific *k*-mer weights are encouraged to be similar to *k*-mer weights in all subclasses the label spans by regularizing on the differences of *k*-mer weights. The overall objective function of SeqUnwinder is: -

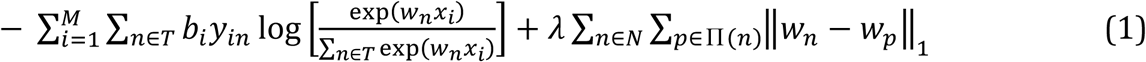

In the above equation; *M* is the total number of genomic loci in all subclasses, *T* is the set of all subclasses, *b*_*i*_ is the weight given to the genomic site *i*, *w*_*n*_ is the *k*-mer weight vector for subclass *n*, *x*_*i*_ is a vector of *k*-mer counts for the genomic site *i*, *y*_*in*_ is a binary indicator variable denoting the subclass of genomic site *i*, *λ* is the regularization co-efficient, Π(*n*) is the set of all labels spanning the subclass *n*, and *w*_*p*_ is the *k*-mer weight vector for label *p*. Values for *bi* are chosen to account for class imbalances. Hence, the value of *b_i_* for a genomic site *i* belonging to class *n* is defined as |*n*_*max*_|/|*n*|, where |*n*| denotes the number of genomic sites in subclass *n* and |*n*_*max*_| denotes the number of genomic sites in the subclass with maximum sites.

### Training the SeqUnwinder model

The *w*_*n*_ and *w*_*p*_ update steps separate out and are iteratively updated until convergence. The *w*_*p*_ update step has a simple closed form solution given by the equation:

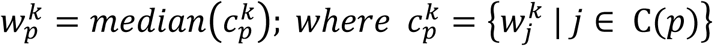

Where 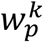 is the *k*^th^ term of the label-*p* weight vector. 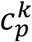 is a set of the *k*^th^ terms of the weight vectors of all the subclasses the label *p* spans.

The *w*_*n*_ update step is: -

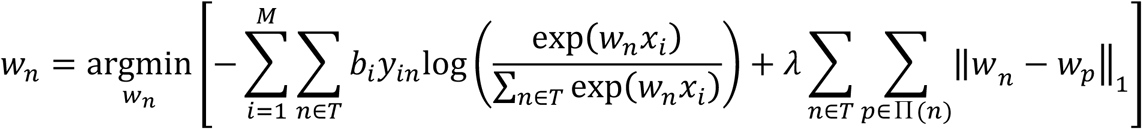

The above equation is solved using the scaled alternating direction method of multipliers (ADMM) framework (Boyd *et al*, 2011). Briefly, the ADMM framework splits the above problem into 2 smaller sub-problems, which are much easier to solve. ADMM introduces an additional variable *z*_*np*_ initialized as follows

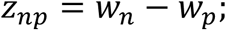

*w*_*x*_ and *z*_*np*_ are iteratively estimated until convergence of the ADMM algorithm.

### Sub-problem 1

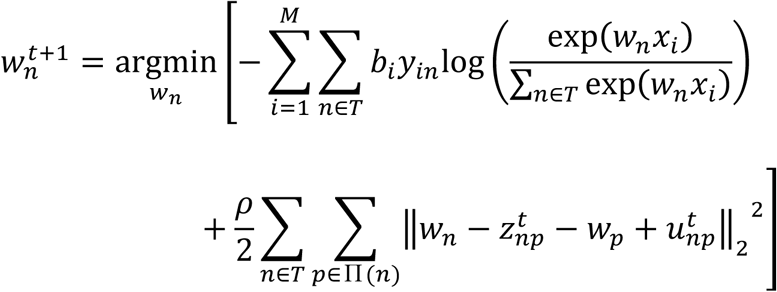

Where *u*_*np*_ is the scaled dual variable. The above sub-problem is solved using the LBFGS (limited-memory Broyden Fletcher Goldfarb Shanno) algorithm (Liu & Nocedal, 1989)

### Sub-problem 2

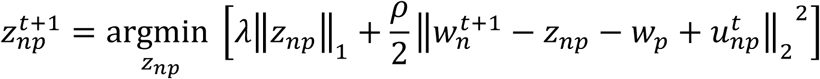

The solution to the above equation is given by the shrinkage function defined as follows: -

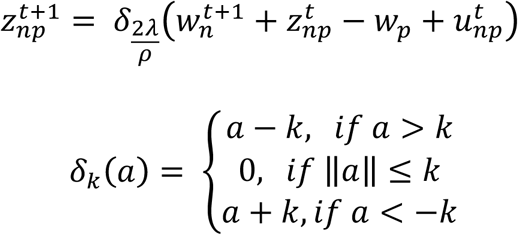

The update step for the scaled dual variable *u*_*np*_ is: -

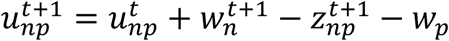

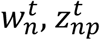, and 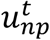are iteratively estimated until convergence. The stopping criteria for the ADMM algorithm is:

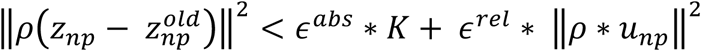

and

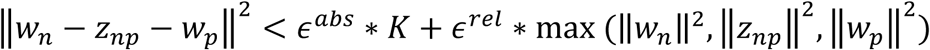

Where *ϵ*^*abs*^ and *ϵ*^*rel*^ are the absolute and relative tolerance, respectively. Of note, to speed up the implementation of SeqUnwinder, a distributed version of ADMM was implemented. Intuitively, the 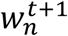 update step is distributed across multiple threads by spitting the M training examples into smaller subsets. The 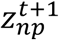 and the 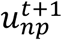 update steps act as pooling steps where the estimates of different threads are averaged. To further speed up convergence, a relaxed version of ADMM was implemented as described in (Boyd *et al*, 2011). In the relaxed version, 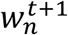 is replaced by 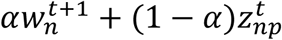 for the 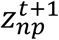 and 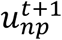 update steps, where α is the over-relaxation parameter and is set to 1.9 as suggested in (Boyd *et al*, 2011)

### Converting weighted k-mer models into interpretable sequence features

While SeqUnwinder models label-specific sequence features using high-dimensional *k*-mer weight vectors, it is often desirable to visualize these sequence features in terms of a collection of interpretable position-specific scoring matrices. To do so, we first scan the *k*-mer models learned during the training process across fixed-sized sequence windows around the input genomic loci to identify local high-scoring regions called “hills”. Label-specific hills are focused regions ranging from 10 to 15bp and are composed of high scoring *k*-mers. We use a threshold of 0.1 to define hills. Next, we cluster the hills using K-means clustering with Euclidean distance metric and *k*-mer counts as features. To speed-up implementation, we restrict the unbiased *k*-mer features to only those *k*-mers that are present in at least 5% of the hills. We use silhouette index (Rousseeuw, 1987) to choose the appropriate value for K. Briefly, we test a range of K values from 2 to 6. For each value of K, we calculate the silhouette index on 30 bootstrap samples. The value of K with highest median silhouette index is chosen as the best value for K. Finally, all clusters with membership less than 10% of the largest cluster are merged by assigning their members to the next closest cluster. MEME (Bailey & Elkan, 1994) is used to identify motifs in different clusters resulting in label specific discriminative motifs. Each *k*-mer model further scores MEME-identified motifs as follows:

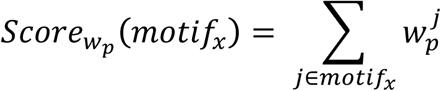

Where *j ∈ motif_x_* is the set of all k-mers that belong to motif *“motif_x_”*.

### Generation of synthetic datasets

To test SeqUnwinder in simulated settings, we generated various synthetic datasets. First, we generated 150bp sequences by sampling from a 2^nd^ order Markov model of the human genome. We then randomly assigned labels to these sequences at different frequencies. The overlap between the labels at the sequences was varied from 0.5 to 0.99. Arbitrarily chosen TF binding motifs were assigned to labels. Each motif instance was sampled from the probability density function defined by the PWM of the motif. Sampled motif instances were inserted at labeled sites at a frequency of 0.7.

### Processing iMN programming data-sets

*Defining early, shared and late binding labels*: MultiGPS was used to call Isl1/Lhx3 binding sites at 12 and 48hrs (datasets were obtained from GSE80321). A q-value cutoff <0.001 was used to call binding sites. All sites with significantly greater Isl1/Lhx3 ChIP enrichment at 12h compared to 48h (q-value cutoff of <0.01) were labeled as “early”. Isl1/Lhx3 binding sites called in both 12 and 48h datasets with a further filter of not being differentially bound (q-value cutoff of <0.01), were assigned as “shared” sites. Finally, all sites with significantly greater Isl1/Lhx3 ChIP enrichment at 48h compared to 12h (q-value cutoff of <0.01) were labeled as “late”. *Defining active and inactive mES annotation labels*: A random forest classifier (see below for implementation details) was trained to classify every Isl1/Lhx3 binding site as either being in accessible/active or inaccessible/unmarked mouse ES chromatin. The classifier was trained using 95 mouse ES ChIP-seq datasets with windowed read-enrichment as predictors. A union list of 1million 500bp regions comprising the enriched domains (see below) of DNaseI, H3K4me2, H3K4me1, H3K27ac, and H3K4me3 was used as the positive set for training the classifier. An equal number of unmarked 500bp regions were randomly selected and used as the negative set for training the classifier. Every binding site that was predicted to be in accessible/active ES chromatin with a probability of greater than 0.6 was placed in the “ES-active” class, while the remaining sites were placed in the “ES-inactive” class.

Enriched domains for DNaseI, H3K4me2, H3K4me1, H3K27ac, and H3K4me3 were identified using the DomainFinder module in SeqCode (https://github.com/seqcode/seqcode-core/blob/master/src/org/seqcode/projects/seed/DomainFinder.java). Contiguous 50bp genomic bins with significantly higher read enrichment compared to an input experiment were identified (binomial test, p-value < 0.01). Further, contiguous blocks within 200bp were stitched together to call enriched domains

Weka’s implementation of Random Forests was used to train the classifier (https://github.com/seqcode/seqcode-core/blob/master/src/org/seqcode/ml/classification/BaggedRandomForest.java). Briefly, the forest contained 10000 trees. Each tree was trained with 10 randomly sampled features on 1% bootstrapped samples of the entire dataset.

### Processing ENCODE datasets

*TF ChIP-seq datasets*: We analyzed 17 TF ChIP-seq ENCODE datasets in three primary cell-lines (GM12878, K562, and H1-hESC). The binding profiles for the factors were profiled using MultiGPS (Mahony *et al*, 2014). All called binding events for TFs were required to have significant enrichment over corresponding input samples (q-value <0.01) as assessed using MultiGPS’ internal binomial test. For a site to be labeled as “shared”, the binding site was required to be called in all the 3 cell-lines. Further, binding sites showing significantly differential binding in any of the possible 3 pair-wise comparisons were removed from the shared set. Binding sites labeled as cell-type specific sites were required to have significantly higher ChIP enrichment compared to other cell-lines. All TF binding sites within 5Kbp of a known TSS (defined using UCSC hg19 gene annotations) were labeled as “promoter proximal” while all sites that were more than 5Kbp from known TSSs were labeled as “distal”.

*DNase-seq datasets*: We analyzed the DHS sites at 6 different tier 1 and 2 ENCODE cell-lines (GM12878, K562, H1-hESC, HeLa-S3, HepG2, HUVEC). The DHS sites were called using in-house scripts. Briefly, contiguous 50bp genomic bins with significantly higher read enrichment compared to an input experiment were identified (binomial test, p-value < 0.01). Further, contiguous blocks within 200bp were stitched together to call enriched domains. A 150bp window around the maximum point of read density at enriched domains was considered as the DHS.

### Annotation of de novo identified motifs

All *de novo* motifs identified using SeqUnwinder were annotated using the cis-bp database. Briefly, *de novo* motifs were matched against the cis-bp database using the STAMP software (Mahony & Benos, 2007). The best matching hit with a p-value of less than 10e-6 was used to name the *de novo* identified motifs.

## Acknowledgments

This work was supported by R01HD079682 NICHD (to EOM). The authors thank Dr. Frank Pugh, Dr. Ross Hardison, and members of the Center for Eukaryotic Gene Regulation at Penn State for helpful discussions.

## Author Contributions

AK and SM conceived the study. AK designed and implemented the SeqUnwinder method and performed all analyses. SV performed iMN ChIP-seq experiments. SM and EOM supervised the work. AK and SM wrote the manuscript.

## Figure Legends

**Figure S1.**
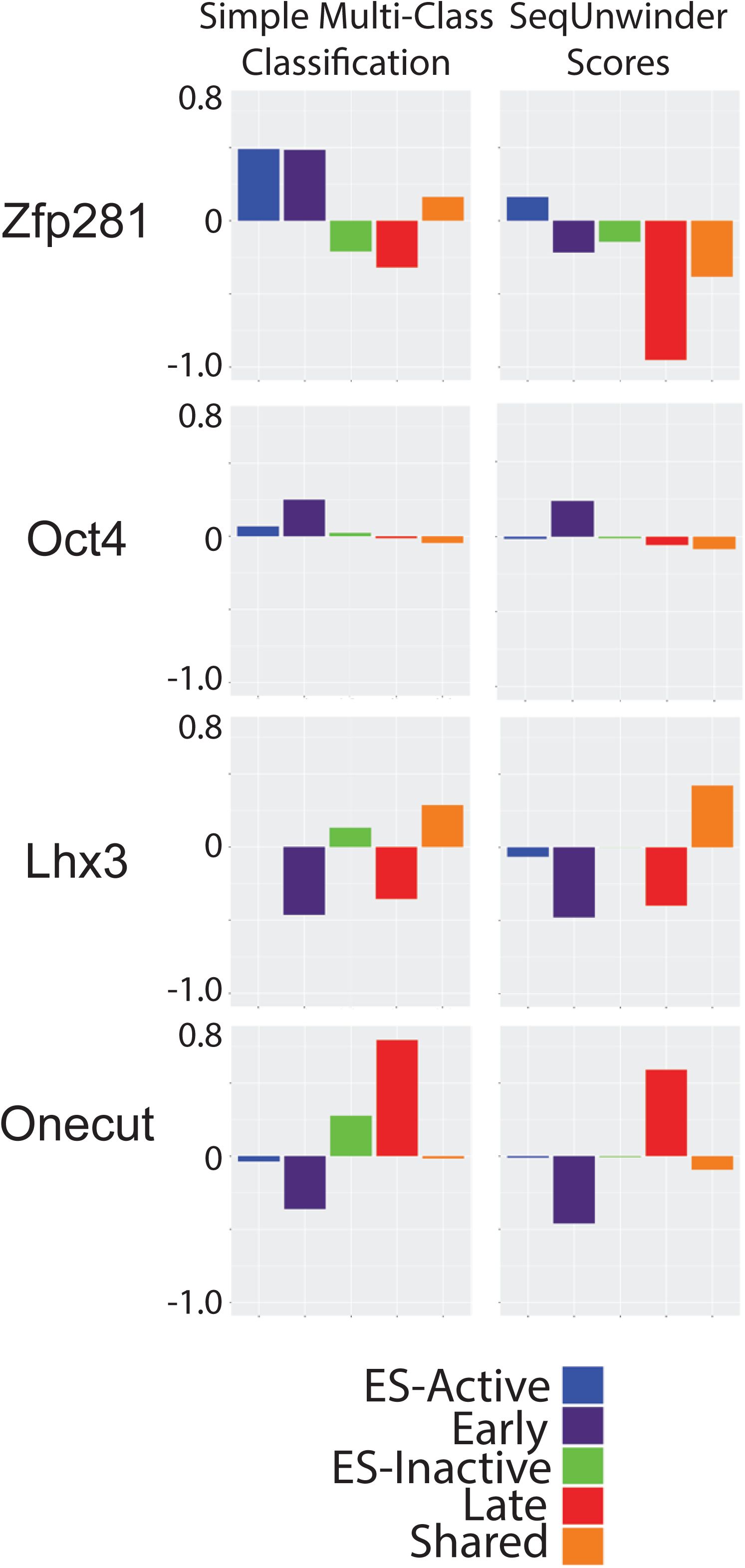
SeqUnwinder and MCC model scores of *de novo* discovered Zfp281-like, Oct4-like, Lhx3-like and Onecut-like motifs.

**Figure S2.**
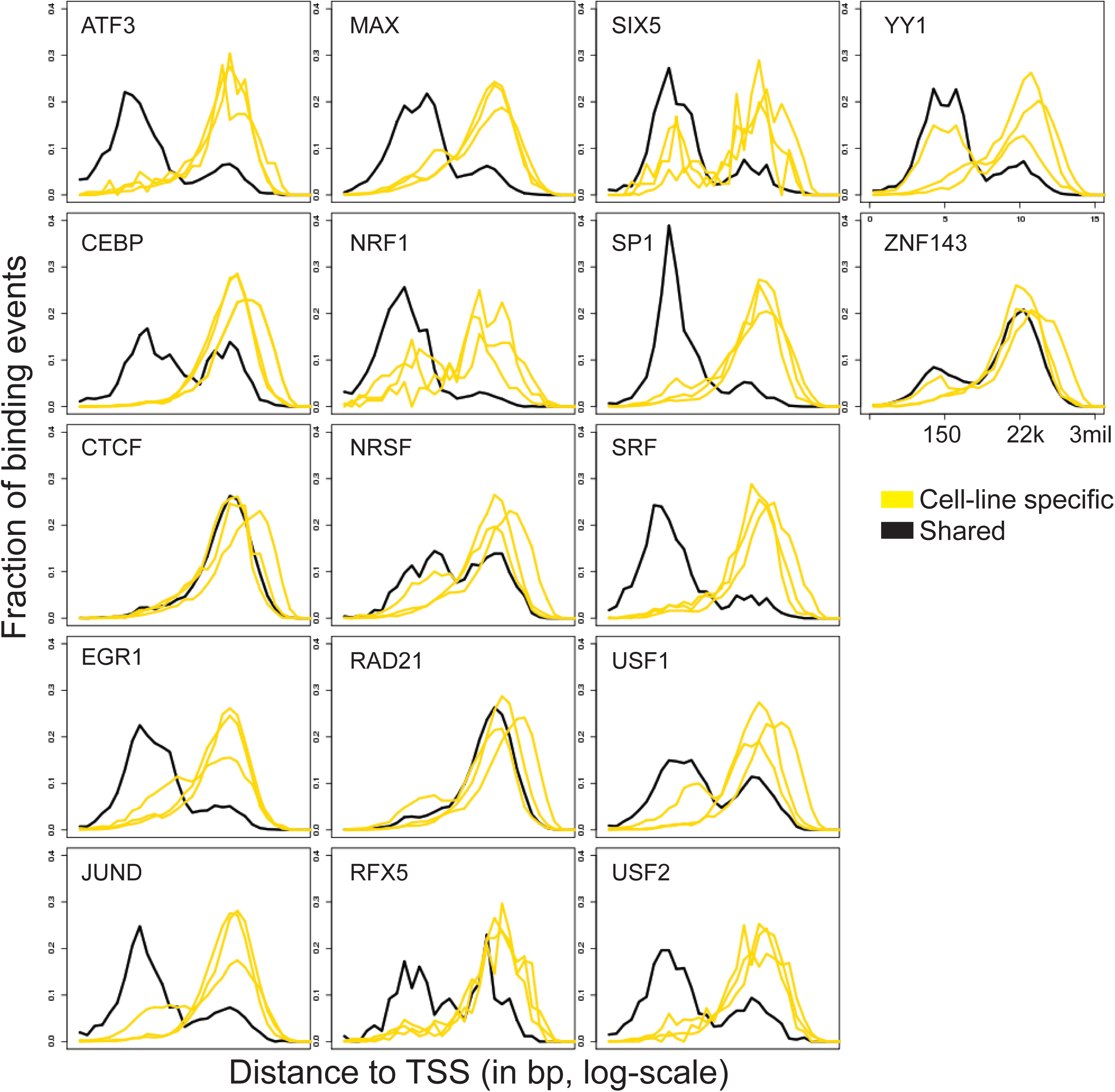
Distance of TF binding events from annotated mRNA TSS for all 17 examined ENCODE TFs, stratified based on “shared” (black) or “cell line-specific” (yellow) labels. The X-axis represents the distance in “bp” in log-scale (natural logarithm).

**Figure S3.**
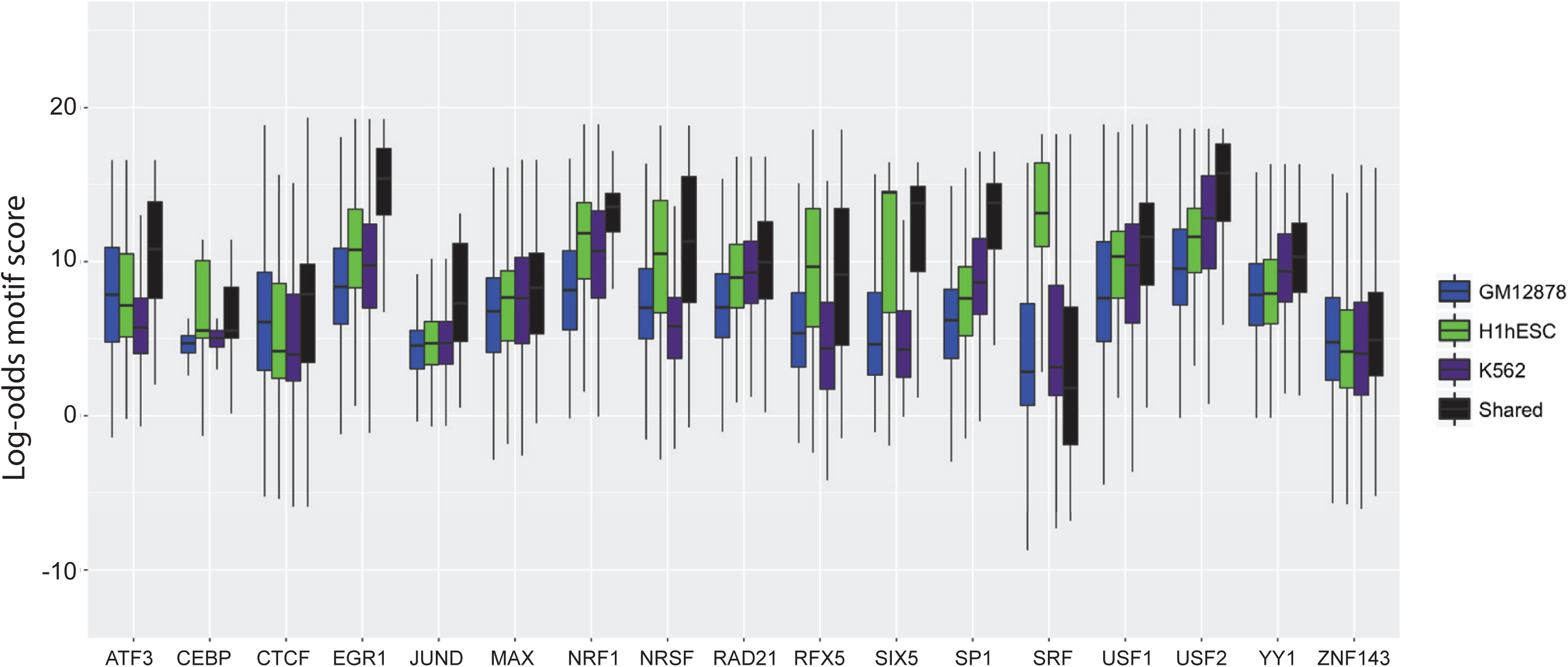
Log-odds score distribution of *de novo* discovered cognate motifs at “shared”, “K562”, “GM12878” and “H1hESC” labeled sites.

